# ALCOHOL AUGMENTS SPONTANEOUS GABAERGIC TRANSMISSION AND ACTION POTENTIAL FIRING IN IMMATURE MURINE CEREBELLAR GOLGI CELLS, LEADING TO ENHANCED INHIBITORY INPUT ONTO CEREBELLAR GRANULE CELLS

**DOI:** 10.1101/2025.09.03.674091

**Authors:** Karick Jotty-Arroyo, Andrea Iturralde-Carrillo, Michael Paffett, Rafael Mancero-Montalvo, C. Fernando Valenzuela

## Abstract

Golgi cells (GoCs) are cerebellar inhibitory interneurons that provide both phasic and tonic GABAergic input to cerebellar granule cells. They receive inhibitory control from Lugaro cells, other GoCs, and cerebellar nuclear inhibitory neurons via GABAergic and glycinergic inputs. Although fetal alcohol exposure is known to impair cerebellar function, its impact on developing GoC physiology remains unclear. We investigated the acute effects of ethanol on GABA_A_ receptor–mediated transmission in GoCs during the mouse equivalent of the human third trimester, a critical window for inhibitory circuit formation. To identify GoCs, we used VGAT-Venus transgenic mice, in which the vesicular GABA transporter promoter drives expression of the Venus fluorescent protein. Whole-cell patch-clamp and loose-patch recordings from postnatal day (P) 6–10 mice revealed that ethanol exposure dose-dependently increased the frequency of action potential–dependent GABA_A_ receptor-mediated spontaneous postsynaptic currents (sPSCs) in GoCs. While ethanol produced variable effects on GoC firing rates, it more consistently enhanced GABA_A_-sPSC frequency in granule cells. We also examined expression of the K^+^–Cl^-^ cotransporter 2 (KCC2), a chloride exporter whose developmental upregulation drives the shift in GABA_A_ receptor signaling from excitatory to inhibitory. Immunohistochemical analysis at P6 showed that GoCs express low levels of KCC2, suggesting that GABA_A_ receptor– mediated currents may remain depolarizing in a subset of GoCs. This property could contribute to ethanol-induced disruption of cerebellar circuit maturation. Together, these findings provide new insight into the cellular mechanisms by which ethanol perturbs inhibitory circuit development in the cerebellum.

## Introduction

Golgi cells (GoCs) are inhibitory neurons with round or polygonal shapes located throughout the granular layer, especially near Purkinje cells [1](Fig. 1). Morphologically, GoCs have a large, irregular soma (10–20 µm), a pale eccentric nucleus with a prominent nucleolus, and a thick axon that bifurcates early and gives rise to numerous collaterals. Their apical dendrites extend into the molecular layer, while basal dendrites remain within the granular layer. Neurochemically, approximately 80% of Golgi cells are both GABAergic and glycinergic, 15% are purely GABAergic, and 5% are exclusively glycinergic [1, 2]. Their axons form part of the cerebellar glomerulus, where GoC axons and mossy fiber terminals converge onto granule cell (GrC) dendrites (Fig. 1) [1, 3]. GoCs provide both phasic and tonic GABAergic inhibition to GrCs, playing a key role in gating information flow into the cerebellar cortex [2]. Additionally, GoCs are involved in regulating long-term synaptic plasticity along the mossy fiber pathway [2]. These neurons are intrinsic pacemakers that fire autonomously at 1-10 Hz [2]. They receive excitatory synaptic inputs from mossy fibers and parallel fibers [2], as well as spillover-mediated glutamatergic transmission from climbing fibers [4]. GoCs have been shown to receive GABAergic and glycinergic inhibition from Lugaro cells [5, 6] and other GoCs [7–9]. Additionally, studies indicate that a significant portion of the inhibitory input to GoCs is provided by cerebellar nuclear inhibitory cells [10, 11].

**Figure 1.**
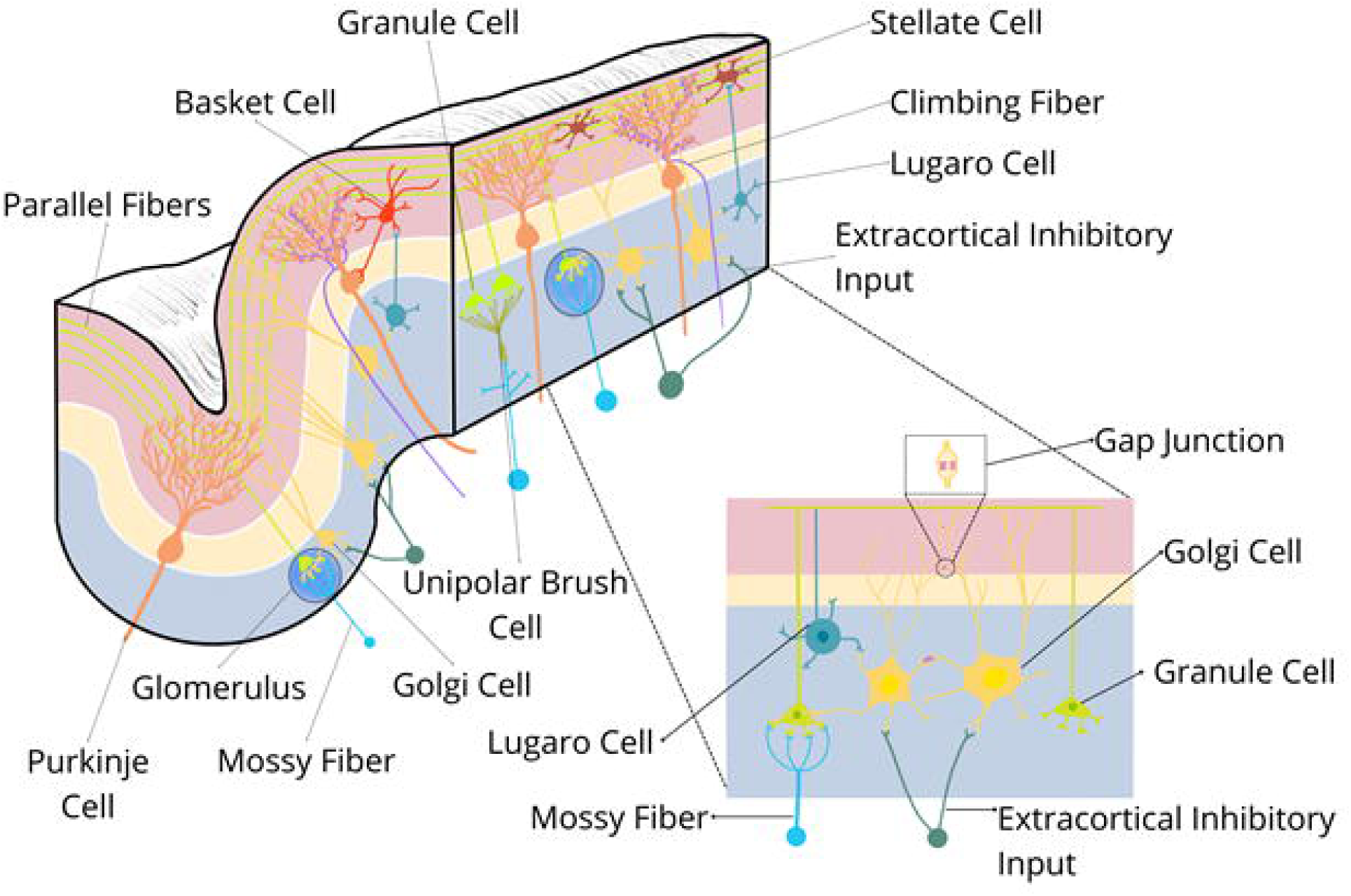
Schematic representation of the cerebellar cortex architecture showing the main neuronal types and their connections. The schematic depicts Purkinje cells, granule cells (GrCs), Golgi cells (GoCs), stellate cells, basket cells, Lugaro cells, and unipolar brush cells, along with parallel, climbing, and mossy fibers, as well as extracortical inhibitory inputs. The cerebellar glomerulus is highlighted as a synaptic unit within the granule cell layer. The inset provides a detailed view of GoCs interconnected via gap junctions, illustrating their excitatory inputs provided by mossy and parallel fibers. The inset also shows GoCs inhibitory inputs provided by Lugaro cells, other GoCs, and extracortical inhibitory inputs, and GrCs.

Developmentally, GoCs originate from the ventricular zone in the rostral-most region of the cerebellar anlage [12]. They mainly undergo radial migration but may also exhibit tangential movements to reach their final positions during cerebellar formation [13]. During migration, GoCs experience terminal mitosis, occurring during late fetal and early postnatal development [12] (Fig. 2). In rodents, at postnatal day 0 (P0), GoCs are predominantly located in the white matter and beneath the Purkinje cell layer [14]. By P4, they are found not only in the white matter but also within the cerebellar cortex, beginning differentiation in the internal GrC layer [14]. By P7, GoCs display their characteristic morphological features, and by P10, they are well differentiated with evident axonal arborizations [14]. Between P15 and P20, GoCs exhibit the typical structure of adult cells and become less abundant than in juvenile mice [14]. Notably, two subpopulations of GoCs have been identified: type 1 GoCs express the gene Gjd2, which encodes dominant gap junctions, as well as the gene Sorcs3, encoding a type-I receptor transmembrane protein that is a member of the vacuolar protein sorting 10 receptor family [15]. In contrast, type 2 GoCs lack Gjd2 expression but display high levels of Nxph1, which encodes neurexophilin-1, a protein that forms a tight complex with α-neurexins, thereby promoting adhesion between dendrites and axons [15]. These findings are consistent with studies indicating that GABAergic GoCs undergo rapid differentiation shortly after birth, especially during the first postnatal week [12]. Despite this growing understanding of GoC subtypes and their development, the precise timeline and mechanisms of GoC synapse formation remain poorly understood.

**Figure 2.**
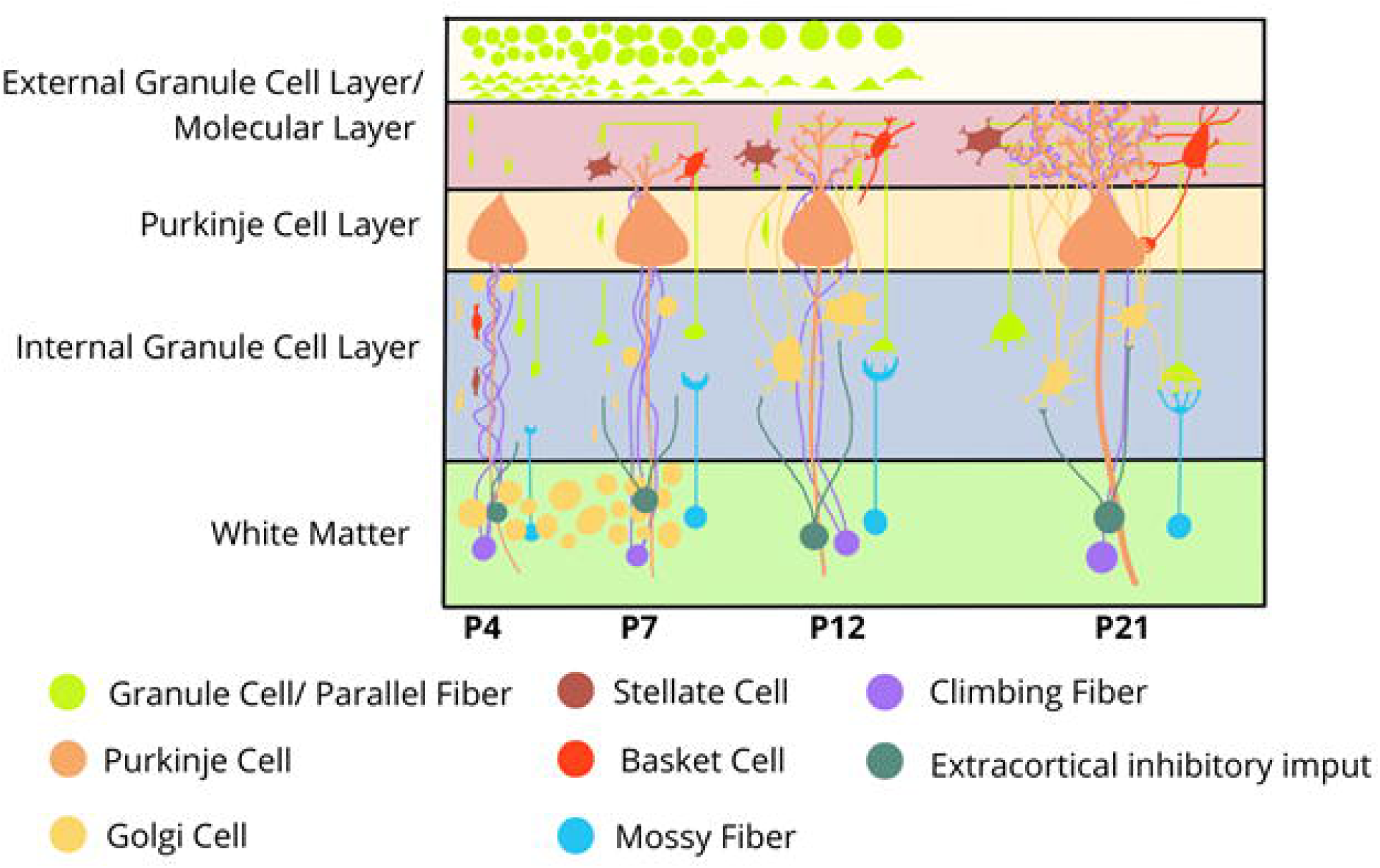
Schematic of cerebellar cortex layers and neuronal organization during development. The diagram depicts the main cellular components across the four layers of the developing cerebellar cortex—external granule cell layer, molecular layer, Purkinje cell layer, and internal granule cell layer—with the white matter also indicated. Neuronal types are color-coded: granule cells (GrCs) and parallel fibers (light green), Purkinje cells (orange), Golgi cells (GoCs; yellow), stellate cells (brown), basket cells (red), climbing fibers (purple), mossy fibers (light blue), and extracortical inhibitory inputs (dark green). From left to right, the sequence illustrates postnatal day (P) 4 to P21. At P4, GoCs are located in the white matter and adjacent to Purkinje cells. By P7, GoCs appear within the internal GrC layer, with slightly enlarged somata and early dendritic extensions but limited arborization. At P12, GoCs develop more elaborate dendritic trees projecting toward the molecular layer, forming expanded contacts with GrCs, mossy fibers, and other GoCs. By P21, GoCs reach their mature morphology, with fully developed dendrites spanning the granular and molecular layers, integrated into the mature cerebellar microcircuit alongside other interneurons.

During early brain development, GABA acts as an excitatory neurotransmitter rather than an inhibitory neurotransmitter due to the high intracellular Cl⁻ concentration in immature neurons. This is driven by the activity of the Na⁺–K⁺–2Cl⁻ cotransporter 1 (NKCC1), which imports chloride into the cell. Under these conditions, activation of GABA_A_ receptors leads to chloride efflux, membrane depolarization, and activation of voltage-gated Ca²⁺ channels and NMDA receptors, ultimately increasing intracellular Ca²⁺ levels. This temporary excitatory role of GABA is essential for key developmental processes, such as neuronal proliferation, migration, and neurite outgrowth [16, 17]. As development advances, the K⁺–Cl⁻ cotransporter 2 (KCC2) becomes upregulated, lowering intracellular chloride levels and shifting GABAergic signaling from excitatory to inhibitory [14, 16–18]. The developmental timeline of this excitatory-to-inhibitory switch in GoCs has not been characterized.

Fetal alcohol exposure is linked to persistent neurological deficits, including motor and cognitive impairments that stem, in part, from cerebellar dysfunction. However, the effects of developmental ethanol exposure on GoC function remain poorly characterized. In juvenile rats, repeated intermittent vaporized ethanol exposure during the rodent equivalent of the third trimester did not significantly alter GoC-mediated tonic or phasic GABA_A_ receptor signaling onto GrCs [19]. Exposure during the same developmental window likewise left the spontaneous activity of mature GoCs and GrCs unchanged, despite reducing the proportion of Purkinje neurons exhibiting complex spike bursts [20]. By contrast, the acute effects of ethanol on mature GoCs are better understood: multiple studies, including our own, have shown that ethanol increases spontaneous GoC firing in both rats and mice [21–28]. In mature GoCs, this effect involves inhibition of neuronal nitric oxide synthase and its downstream targets—the Na^+^/K^+^-ATPase, a key regulator of intrinsic pacemaker activity—and suppression of a quinidine-sensitive K^+^ channel [21, 25, 26]. Whether ethanol similarly affects the excitability of developing GoCs or modifies their synaptic inputs, however, is unknown. To address this gap, we examined the acute effects of ethanol on GABA_A_ receptor-mediated transmission in GoCs during the mouse equivalent of the third trimester of human pregnancy, a critical window for establishing inhibitory circuitry in the cerebellar cortex.

## Materials and methods

### Animals

Vesicular GABA transporter (VGAT)-Venus transgenic mice, which express fluorescently labeled GABAergic and glycinergic neurons throughout the brain, were generously provided by Dr. Yanawaga (Gunma University, Japan) [29]. Venus is a yellow fluorescent protein variant characterized by efficient maturation and enhanced resistance to the intracellular environment [30]. VGAT-Venus mice were maintained as heterozygotes. Female VGAT-Venus or wild-type C57BL/6 mice were mated with either VGAT-Venus or wild-type C57BL/6 males, respectively. Males were removed once pregnancy was confirmed. After birth, postnatal day (P)1–P2 pups were screened for Venus expression using a “miner’s lamp” emitting 460–495 nm light; yellow fluorescence in the brain was detected using a 520–550 nm filter (Biological Laboratory Equipment Maintenance and Service LTD, Budapest, Hungary).

### Immunohistochemistry

Three P6 VGAT-Venus mice were used for these experiments; the mice were randomly chosen and their sex was not determined. Mice were injected intraperitoneally with ketamine and xylazine (250 mg/kg and 25 mg/kg, respectively). Mice were transcardially perfused with 32 °C phosphate-buffered saline (PBS, Thermo Fisher Scientific pH 7.4) containing 1 g/L procaine hydrochloride (Sigma-Aldrich, St. Louis, MO) and 1 USP units/mL heparin (Sagent Pharmaceuticals, Shaumburg, IL) for 2 minutes, followed by room-temperature 4% paraformaldehyde (PFA, Sigma-Aldrich) in PBS for an additional 2 minutes, and then 5 minutes of ice-cold 4% PFA in PBS. After extraction, the brains were stored in 4% PFA for 48 hours on a rotating shaker at 4 °C, and then transferred to 30% sucrose (w/v in PBS) for another 48 hours. Cerebella were embedded in Tissue-Tek^®^ O.C.T. Compound (Sakura Finetek, Torrance, CA) on disposable specimen molds, frozen on dry ice, and stored at −80 °C. Before sectioning, the cerebella were allowed to equilibrate at −20 °C for 3 hr. Parasagittal sections (50 μm-thick) containing the cerebellar vermis were sectioned using a cryostat (Model 505E, Microm Gmb, Walldorf, Germany) and stored at −20 °C in multiple-well plates with a cryoprotectant solution (0.1M phosphate buffer pH 7.4, 25% glycerol, 25% ethylene glycol).

The staining procedure was performed over two days. On Day 1, the floating sections were washed three times with 500 μl of 1X PBS and incubated in a blocking solution containing 1% bovine serum albumin (Catalog # A4737, Sigma Aldrich), 5% donkey serum (Catalog # 017000121, Jackson Immuno Research, West Grove, PA), and 0.2% Triton X-100 (Sigma Aldrich). The sections were then incubated with 300 μl of primary antibody solution (1:1000 of rabbit anti-K^+^-Cl^−^ cotransporter (KCC2) antibody (Catalog #: C2366, Sigma Aldrich) for 18 hours with gentle shaking at 4 °C. On Day 2, the sections were washed with 500 μl of 1X PBS and incubated for 30 minutes at room temperature in a blocking solution containing 1% bovine serum albumin and 5% donkey serum (Catalog # 017000121, Jackson Immuno Research). They were then incubated for 2 hours at room temperature with 300 μl of a secondary antibody solution (1:1000 Alexa Fluor 555, donkey anti-rabbit IgG, Catalog # A-31572, Thermo Fisher Scientific, Waltham, MA). During the last 20 minutes of the previous incubation, nuclei staining was performed with 4’,6-diamidino-2-phenylindole (DAPI, dilution 1:500, Thermo Fisher Scientific). The sections were rinsed three times with 500 μl of 1X PBS, mounted on Superfrost Plus microscope slides (Thermo Fisher Scientific) using Fluoromount G media (Southern Biotech, Birmingham, AL), covered with glass coverslips, and stored at 4 °C until imaging acquisition.

### Microscopy

Fluorescence imaging was done using a Zeiss Axioscan Z1 (Preclinical Core, Center for Brain Recovery and Repair, UNM-HSC) at 20X magnification. Images of the cerebellar vermis were captured across eight brain sections per animal and then averaged. Three fluorescence channels were analyzed: blue for DAPI nuclear labeling, green for Venus-positive interneurons, and orange for KCC2-positive cells. Two investigators blinded to the experimental conditions independently conducted image analysis using Fiji (ImageJ software) and QuPath [31, 32]. The external granule cell, Purkinje, and granular layers were outlined, and KCC2 fluorescence intensities were quantified. Results from both investigators were averaged for consistency.

To extend these analyses, a subset of the sections was visualized by spinning disk confocal microscopy (Evident IX83/Yokogawa CSU-W1, Tokyo, Japan). Cerebellar macro-resolution acquisition was performed using a 30X/1.05NA silicone oil immersion objective with standard disk pinhole size (50 µm) and utilizing Evident cellSens Dimension software to stitch a tiled montage of intact cerebellar sections from P6 mice. A 3-color sequential scanning strategy was deployed between DAPI, Venus, and Alexa Fluor 555 where fluorophores were excited at 405nm, 488nm and 561nm, respectively. Multi-channel images were acquired using a FusionBT sCMOS camera (Hamamatsu Photonics, Shizuoka, Japan), and focus mapping of non-planar lobe sections was applied to ensure a consistent focal plane before acquiring individual images that comprised the 2D cerebellar montage. Additional high-resolution 3D images of Venus-positive Golgi cells located in the granular layer were assessed along with KCC2-positive cells using a 60X/1.3NA silicone immersion oil objective. All 2D and 3D images were deconvolved as previously described [33, 34] before colocalization analysis using Huygens Essential software (Scientific Volume Imaging, Hilversum, Netherlands).

### Electrophysiology

P6-10 VGAT-Venus mice were euthanized by rapid decapitation under anesthesia with ketamine (250 mg/kg I.P.). Brains were extracted and incubated for 2 minutes in cold sucrose artificial cerebrospinal fluid (aCSF) containing (in mM): 220 Sucrose, 2 KCl, 1.25 NaH_2_PO_4_, 26 NaHCO3, 12 MgSO_4_, 10 Glucose, 0.2 CaCl_2_, and 0.43 ketamine, pre-equilibrated with 95% O_2_/5% CO_2._ The cerebellum was embedded in 4% low-melting point agarose at 37 °C in culture dishes and then placed on ice. Once the agarose hardened, the vermis of the cerebellum was sliced parasagittally in cold sucrose aCSF at 200 µm using a vibrating tissue slicer (Leica Microsystems, Bannockburn, IL). Slices were allowed to recover for 40 minutes at 36 °C, transferred to a chamber filled with normal aCSF containing (in mM): 126 NaCl, 2 KCl, 1.25 NaH_2_PO_4_, 26 NaHCO_3_, 10 Glucose, 1 MgSO_4_, 2 CaCl_2,_ and 0.4 ascorbic acid, and were continuously bubbled with 95 % O_2_/5 % CO_2_. Slices were stored in normal aCSF at 21-22°C.

Whole-cell patch-clamp recordings were performed in a chamber perfused with aCSF at a rate of 2-3 ml/min and maintained at 32-33°C. Neurons were visualized using infrared-differential interference contrast microscopy and recordings were performed with a Multi-Clamp 700B amplifier (Molecular Devices, Sunnyvale, CA). GoCs and GrCs were identified as Venus-positive and Venus-negative cells in the GrC layer. Patch pipettes (tip resistance = 3-5 MΩ) were filled with an internal solution containing (in mM): 140 CsCl, 10 HEPES (pH 7.3), 1 EGTA, 4 magnesium ATP, 0.4 GTP, and 4 QX-314 (Tocris-Cookson, Ellisville, MO), pH 7.25, osmolarity 280-290 mOsm. GABAergic transmission was isolated by blocking AMPA and NMDA receptors using kynurenic acid (3 mM). The holding potential was −70 mV. Loose-patch cell-attached recordings were performed as previously described [26]. Electrophysiological recordings were analyzed with Clampfit-10 (Molecular Devices) or MiniAnalysis (Synaptosoft, Decatur, GA).

### Statistical Analysis

Statistical analyses were conducted using Prism (Version 10, GraphPad Software, San Diego, CA). The ROUT test was used to detect outliers (Q = 1%). Data were analyzed by the Shapiro-Wilk normality and lognormality tests. Data that followed a normal or lognormal distribution (transformed to log scale in the latter case) were analyzed using mixed-level ANOVA. Data that did not follow a normal distribution were analyzed using the Kruskal-Wallis test. Planned comparisons (with respect to the time point preceding alcohol application) were done with Dunnett’s test corrected for multiple comparisons. Effect sizes for statistically significant planned comparisons were calculated using Cohen’s d (using https://lbecker.uccs.edu/). As a reference, values of 0.2, 0.5, and >0.8 indicate small, medium, and large effect sizes, respectively. An alpha level of 0.05 was used for all statistical tests. Data are presented as mean ± SEM. For the immunohistochemistry studies, the unit of determination was an animal. For electrophysiological experiments, the unit of analysis was the cell, as the within-cell design enabled direct comparison across conditions (baseline, acute ethanol exposure, and washout). Detailed results of the analyses, including complete statistical notation, are provided in the Supplementary Data Tables.

## Results

### Developing GoCs express low KCC2 levels

Using immunohistochemistry and fluorescent microscopy with a slide scanner, we examined the expression of the Cl^-^ exporter KCC2 in the cerebellar vermis of P6 VGAT-Venus mice. KCC2 plays a critical role in determining whether GABA_A_ receptors function as excitatory or inhibitory ligand-gated ion channels. KCC2 expression was highest in the Purkinje cell layer, followed by the GrC layer and the external granule cell/molecular layer (Fig. 3). As shown in Figure 3B, KCC2 was concentrated around Purkinje cell somata, whereas expression in the GrC and molecular layers appeared more diffuse. Fig. 3D shows the average results obtained from 3 mice. One-way ANOVA yielded a p = 0.0002. The results of Tukey’s post hoc tests were: PL vs. EGL/ML, p=0.0002, Cohen’s *d* = 6.714; PL vs. IGL, p=0.034, Cohen’s *d* = 1.86; IGL vs. EGL/ML, p=0.0018, Cohen’s *d* = 12.54.

**Figure 3.**
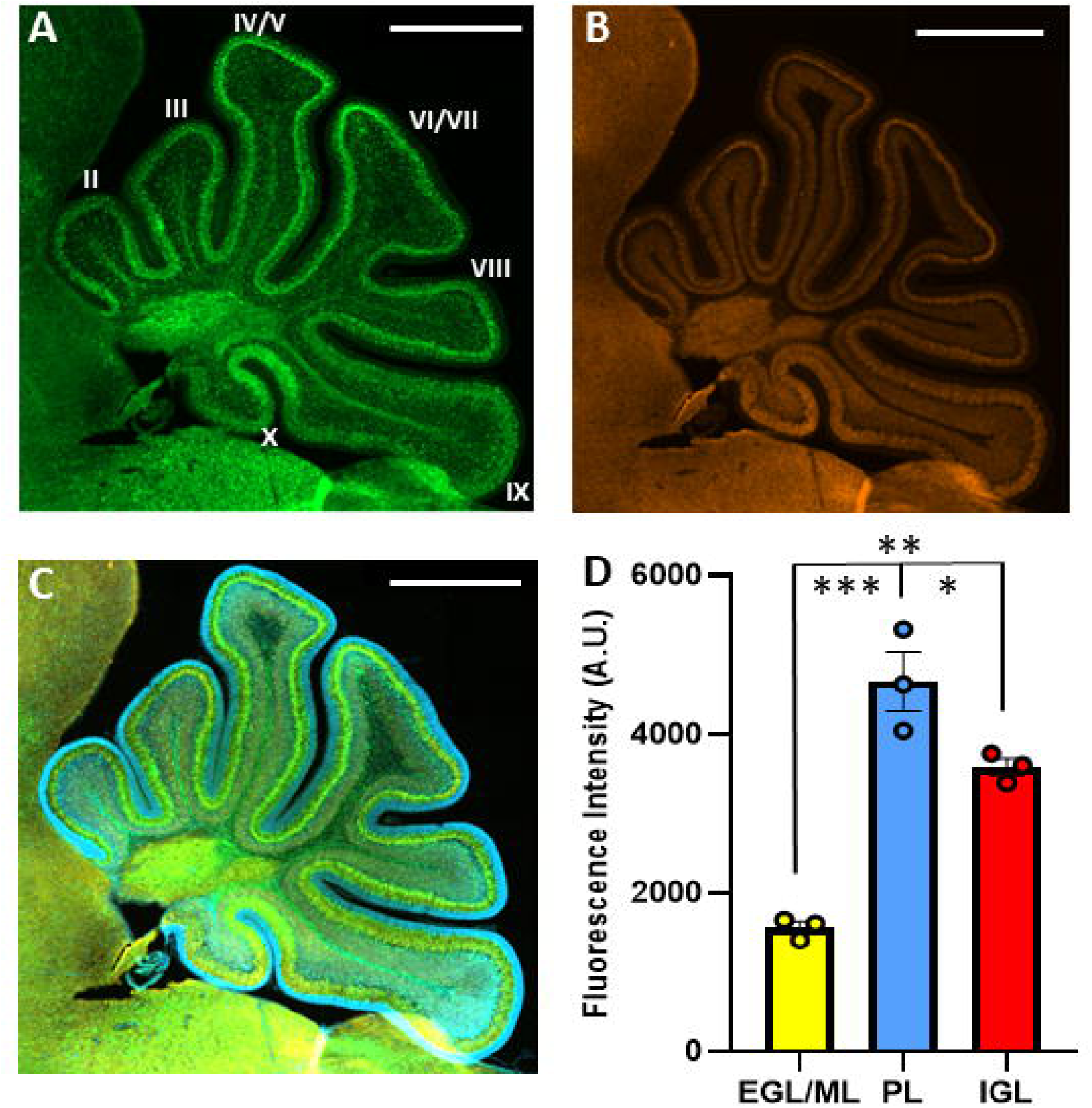
Layer-specific expression of KCC2 in the cerebellar cortex of postnatal day 6 mice. Representative images of a parasagittal section from the cerebellar vermis of VGAT-Venus mice. Shown are individual fluorescence channels for A) Venus (green), B) KCC2 (orange), as well as a C) merged image that also includes DAPI fluorescence (scale bar: 500LJμm). Images were acquired using a Zeiss Axioscan Z1 slide scanner at 20× magnification. D) Quantification of KCC2 expression across distinct layers of the cerebellar cortex: external granule layer/molecular layer (EGL/ML), Purkinje layer (PL), and internal granule layer (IGL). Data were analyzed by one-way ANOVA, which revealed a p=0.0002 (Tukey’s post hoc test: ***p=0.0002; **p=0.018; *p=0.034; n = 3 pups).

Confocal microscopy was used to determine the neuronal subtypes that expressed KCC2. Direct visual identification of Venus and KCC2 immunolabeling overlap was observed in the Purkinje cell layer of P6 cerebellar sections (Fig 4A-B). In GoCs, Venus and KCC2 signals showed partial colocalization, restricted to specific regions of the cell body, whereas KCC2 expression was more abundant in the surrounding GrCs (Fig. 4C). These results prompted us to pursue colocalization analyses. Regions of interest were carefully traced for each layer using Huygens Colocalization Analyzer. The thresholds for Venus/KCC2 channels were uniformly set to produce a defined overlap mask for positive green-red merged events, visualized as orange to yellow. Pearson’s R ± SEM colocalization values were recorded in all cerebellar vermis lobules (Fig 4D). We found that there was lower Venus/KCC2 co-localization in the external and internal GrC layers at P6 (p<0.0001 by mixed effects ANOVA; Tukey’s post hoc tests: EGL/ML vs. PL, p=0.0039, Cohen’s *d* = 4.416; PL vs. IGL, p=0.0105, Cohen’s *d* = 2.893).

**Figure 4.**
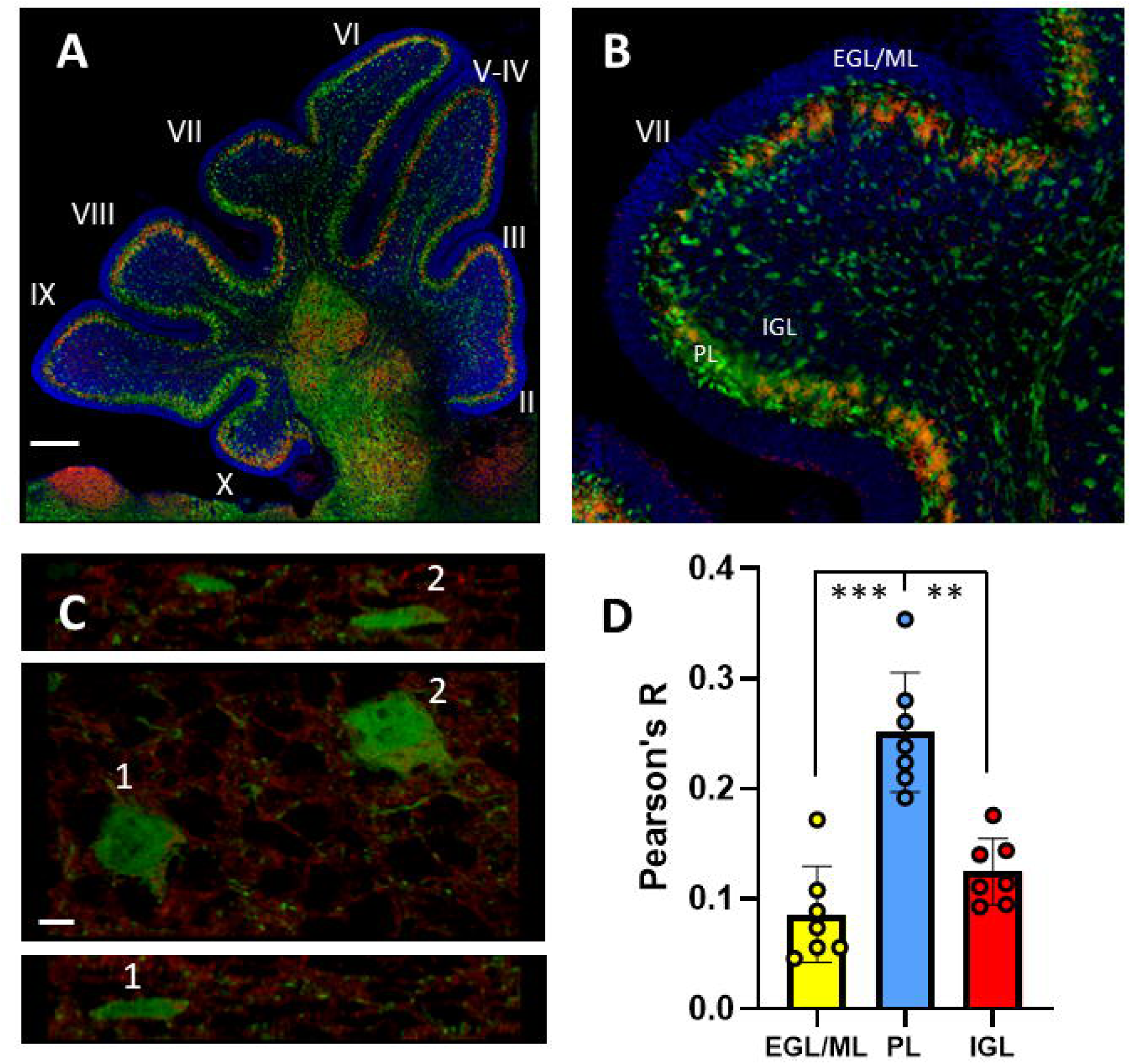
GoCs express low levels of KCC2 at postnatal day 6. A) Three-color confocal montages showing VGAT-Venus and KCC2 immunofluorescence in a cerebellar vermis slice. These images highlight the overlap of Venus and KCC2 signals within specific regions of the Purkinje cell layer, in contrast to the more limited overlap observed in both the external and internal granule cell layers. Images were acquired using an Evident IX83 spinning disk confocal microscope at 20× magnification. B) An enlarged image of lobule VII provides enhanced visualization of these patterns. C) Higher-magnification image and orthogonal projections for GoCs #1 and #2 illustrating the limited co-localization of KCC2 with Venus-positive GoCs. Note the more abundant expression of KCC2 in the surrounding GrCs. Images were also acquired using an Evident IX83 spinning disk confocal microscope but at 60× magnification. D) Summary graph depicting Pearson’s R, an index of colocalization between Venus and KCC2. Each point represents a lobule. External granule cell layer/molecular layer (EGL/ML), Purkinje cell layer (PL), and internal granule cell layer (IGL). (p<0.0001 by mixed effects ANOVA; Tukey’s post hoc test: EGL/ML vs. PL, ***p=0.0039; PL vs. IGL, **p=0.0105).

### Acute ethanol exposure increases GABA_A_-sPSC frequency, but not GABA_A_-mPSC frequency, in immature GoCs

We recorded GABA_A_-sPSCs and GABA_A-_mPSCs from GoCs at P6–10 using the whole-cell patch-clamp configuration. Baseline characteristics of these events are summarized in Table 1. Acute application of ethanol produced a reversible, dose-dependent increase in the frequency of GABA_A_-sPSCs (Fig. 5A, B). Mixed effects ANOVA found a significant effect of time (p=0.0015), and ethanol concentration (p=0.02); planned comparison with Dunnett’s test for 50 mM ethanol: 150 vs. 240 s, p=0.0203, Cohen’s *d*=1.31; 150 vs. 390 s, p=0.0334; Cohen’s *d*=1.11). Mixed effects ANOVA of the amplitude (Fig. 5A, C) data found a significant time x ethanol interaction (p=0.0002) and planned comparison with Dunnett’s test revealed two significant effects for 50 mM ethanol (150 vs. 660 s, p=0.044, Cohen’s *d*=1.18, and 150 vs. 690 s, p=0.025, Cohen’s *d*=1.46). There were no significant effects on area (Fig. 5A, D; Time p=0.066; EtOH Concentration p=0.1848; Time x EtOH Concentration p=0.4227), rise time 10– 90% (Fig. 5A, E; Time p=0.4663; EtOH Concentration p=0.2443; Time x EtOH Concentration p=0.5158), or decay time (Fig. 5A, F; Time p=0.219; EtOH Concentration p=0.433; Time x EtOH Concentration p=0.231). Mixed effects ANOVA of the half-width (Fig.5A, G) found a significant time x ethanol interaction (p=0.0265) but not time (p=0.219) or EtOH concentration (p=0.433); planned comparison with Dunnett’s test did not reveal any significant effects. For comprehensive statistical results and complete statistical notation, refer to the Supplementary Data Tables.

**Figure 5.**
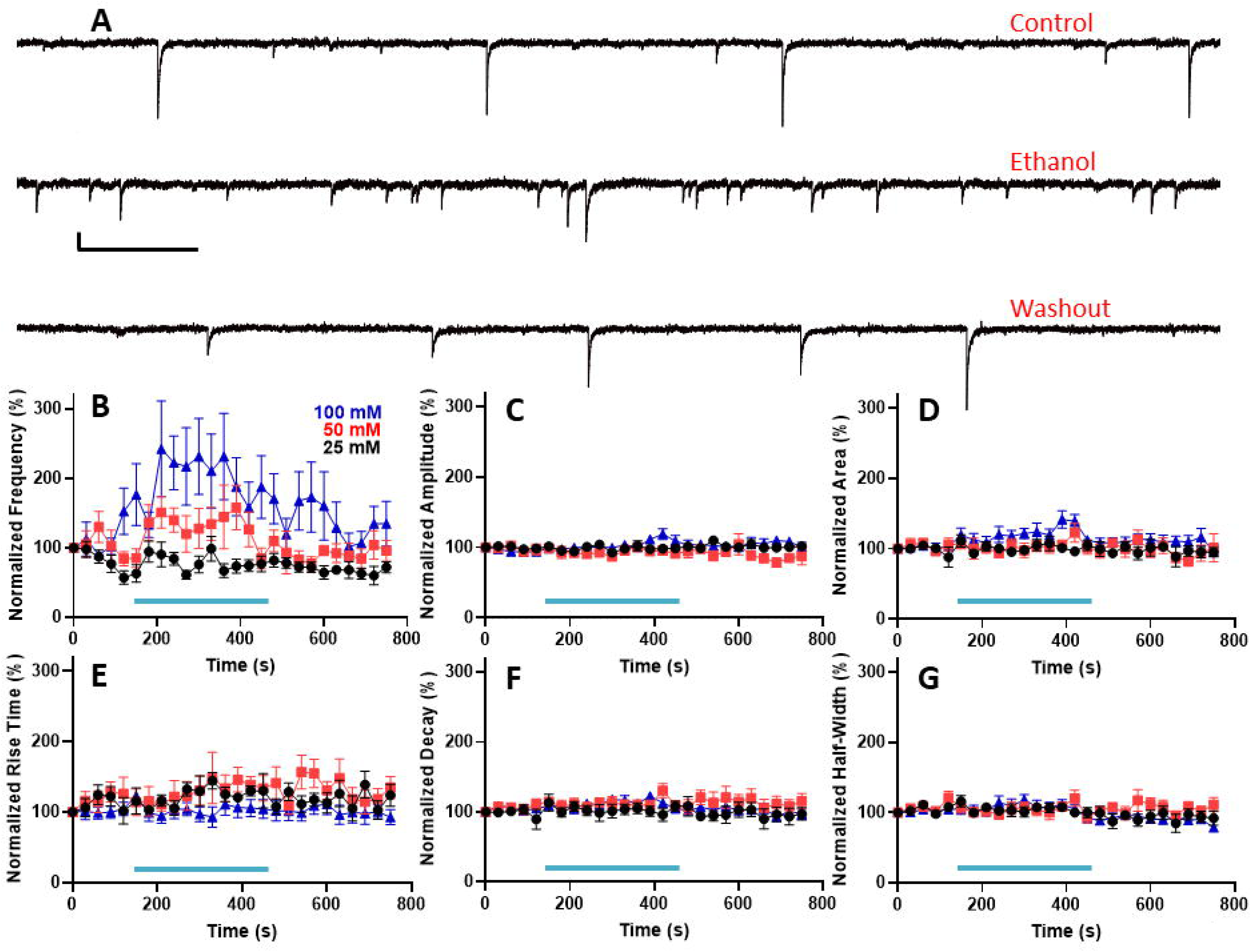
Acute ethanol exposure dose-dependently increases the frequency of action potential-dependent GABA_A_ receptor-mediated spontaneous postsynaptic currents (GABA_A_-sPSCs) in Golgi cells at postnatal days 7-10. A) Representative GABA_A_-sPSC traces from a Golgi cell recording obtained in the absence of ethanol (control), during application of ethanol (50 mM), and during washout. Scale bar is 1.638 s × 80 pA. Mean effect of 25 (n=8), 50 (n=7), and 100 mM (n=9) ethanol on the following GABA_A_-sPSC parameters: B) frequency, C) amplitude, D) area, E) rise time 10-90%, F) decay time, and G) half-width. Data were normalized to the zero-time point. The light blue line represents the timing of ethanol application during the slice electrophysiological recording. Mixed effects ANOVA found significant effects of ethanol on frequency (time, p=0.0015; ethanol concentration, p=0.02; interaction, p=0.1457; planned comparison with Dunnett’s multiple: 150 vs. 240 s, p=0.0203; 150 vs. 390 s, p=0.0334) and amplitude (time, p=0.4; ethanol concentration, p=0.25; interaction, p=0.0002; planned comparison with Dunnett’s multiple: 150 vs. 660 s, p=0.044; 150 vs. 690 s, p=0.025).

**Table 1.** Baseline Postsynaptic Current Parameters (Mean ± SEM)

| **Parameters** | **GoC sPSC (24)** | **GoC mPSC (8)** | **GrC sPSC (n=10)** |
| --- | --- | --- | --- |
| Frequency (Hz) | 1.22 ± 0.25 | 1.56 ± 0.19 | 0.25 ± 0.1 |
| Amplitude (pA) | 70.32 ± 3.72 | 31.5 ± 1.17 | 77.69 ± 12.03 |
| Area (pA/ms) | 829.9 ± 64.3 | 176.7 ± 21.68 | 489.0 ± 126.2 |
| Rise time 10-90% (ms) | 3.09 ± 0.14 | 1.34 ± 0.07 | 1.27 ± 0.12 |
| Decay time (ms) | 14.4 ± 0.72 | 6.66 ± 0.65 | 8.28 ± 0.97 |
| Half-width (ms) | 9.58 ± 0.41 | 4.22 ± 0.38 | 5.88 ± 0.74 |

Recordings of GABA_A-_sPSCs include both action potential–dependent events, driven by spontaneous neuronal firing, and action potential–independent events resulting from the spontaneous fusion of GABA-containing vesicles with the presynaptic membrane. The latter are referred to as GABA_A_-mPSCs. To isolate these miniature events, we assessed the effects of ethanol in the presence of 1 µM tetrodotoxin (TTX), a voltage-gated sodium channel blocker that eliminates action potential–dependent activity. The effects of TTX alone were examined in a separate subset of GoCs from P8 mice not used for ethanol testing. In this group, TTX significantly reduced GABA_A_-sPSC frequency by 40%, from 0.78 ± 0.22 Hz to 0.47 ± 0.18 Hz (n=7; Shapiro-Wilk normality test p = 0.03 for both groups; W=-26; p = 0.03 by Wilcoxon matched-pairs signed rank test; Cohen’s *d* =0.57) and the amplitude of GABA_A_-sPSCs by 13%, from 33.7 ± 1.59 pA to 29.3 ± 1.78 pA (n=; Shapiro-Wilk normality test p=0.94 and 0.14 for the control and TTX groups, respectively; t(6) = 3.115; p = 0.02 by paired t-test; Cohen’s *d*=0.97). Acute application of 50 mM ethanol did not significantly alter the frequency (Fig. 6A, B; p=0.2956), amplitude (Fig. 6A, C; p=0.37), area (Fig. 6A, D; p=0.1405), rise time 10–90% (Fig. 6A, E; p=0.446), decay time (Fig. 6A, F; p=0.1567), or half-width (Fig. 6A, G; p=0.153) of GABA_A_-mPSCs. Statistical details for all comparisons are provided in the Supplementary Data Table, along with complete statistical notation.

**Figure 6.**
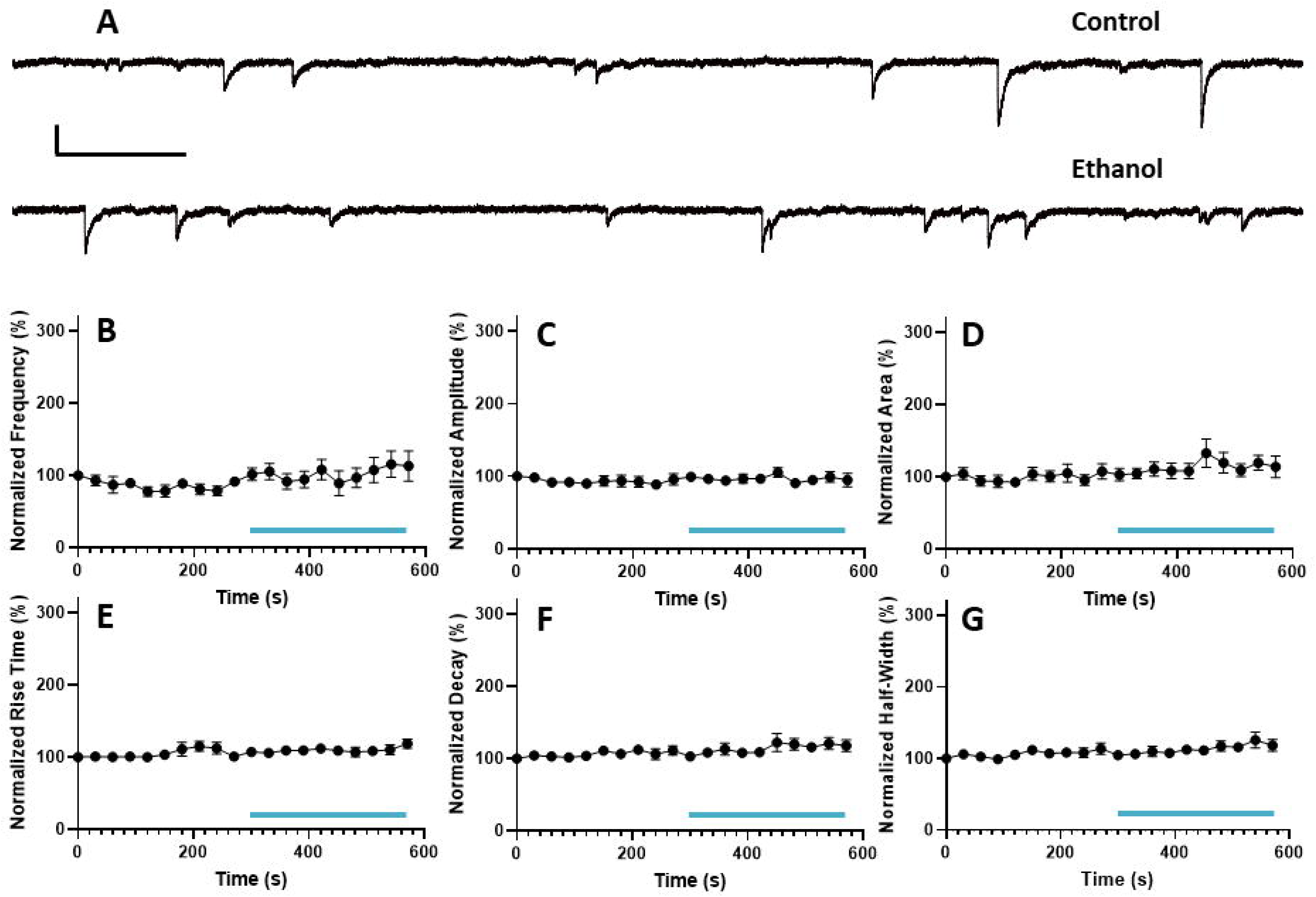
Acute exposure to 50 mM ethanol does not affect action potential-independent GABA_A_ receptor-mediated miniature postsynaptic currents (GABA_A_-mPSCs) in Golgi cells at postnatal days 7-10. A) Representative GABA_A_-mPSC traces from a Golgi cell recording obtained in the absence of ethanol (control), during application of ethanol (50 mM), and during washout. Scale bar is 256 ms × 40 pA. Mean effect of 50 mM ethanol (n=8) on the following GABA_A_-mPSC parameters: B) frequency, C) amplitude, D) area, E) rise time 10-90%, F) decay time, and G) half-width. Data were normalized to the zero-time point. The light blue line represents the timing of ethanol application during the slice electrophysiological recording. Mixed effects ANOVA found no significant effects of ethanol.

### Acute ethanol exposure increases spontaneous action potential firing in developing GoCs and GABA_A_-sPSC frequency in immature GrCs

The ethanol-induced increase in GABA_A_-sPSC frequency may enhance excitatory GABA_A_ receptor signaling in GoCs lacking KCC2 expression, potentially leading to elevated spontaneous action potential firing. This possibility was explored using the loose-patch cell-attached configuration that does not disrupt the composition of the intracellular fluid of GoC. Acute exposure to 50 mM ethanol increased firing frequency in some cells; however, the overall effect was variable and did not reach statistical significance (Fig. 7A–B; *p* = 0.19). In some GoCs, this increase was, at least, reversible (Fig. 7B). Complete statistical values and notation are provided in the Supplementary Data Tables.

**Figure 7.**
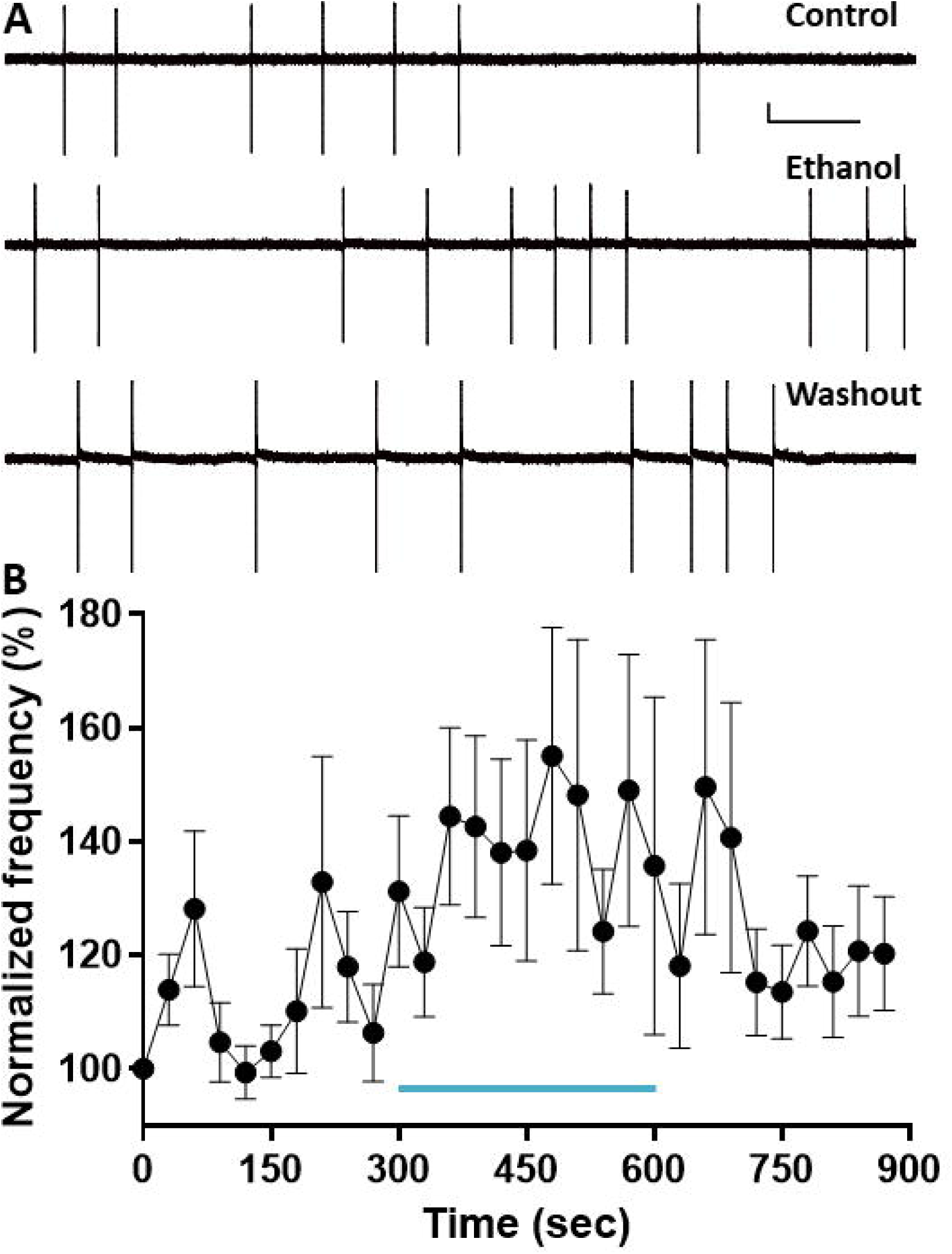
Acute exposure to 50 mM ethanol increases spontaneous action potential firing in Golgi cells at postnatal days 7-10. A) Representative trace of a loose-patch cell-attached recording obtained from a Golgi cell in the absence of ethanol (control), during application of ethanol (50 mM), and during washout. Scale bar is 512 ms × 40 pA. B) Mean effect of 50 mM ethanol (n=10) on spontaneous action potential firing frequency. Data were normalized to the zero-time point. The light blue line represents the timing of ethanol application during the slice electrophysiological recording. Mixed effects ANOVA found no significant effect of ethanol.

We recorded GABA_A_-sPSCs from GrCs at P6–10 using the whole-cell patch-clamp configuration. Baseline characteristics of these events are summarized in Table 1. We investigated whether the ethanol-induced enhancement of GoC firing leads to increased spontaneous, action potential–dependent GABA release at immature GoC–GrC synapses. Mixed effects ANOVA revealed that application of 50 mM ethanol significantly increases the frequency of GABA_A_-sPSCs (Fig. 8A, B; p=0.0094); however, planned comparisons did not reveal any significant effects. There were no significant effects on amplitude (Fig. 8A, C; p=0.37), area (Fig. 8A, D; p=0.954), 10–90% rise time (Fig. 8A, E; p=0.1658), decay time (Fig. 8A, F; p=0.2662), or half-width (Fig. 8A, G; p=0.2964). Full statistical results and notation are provided in the Supplementary Tables.

**Figure 8.**
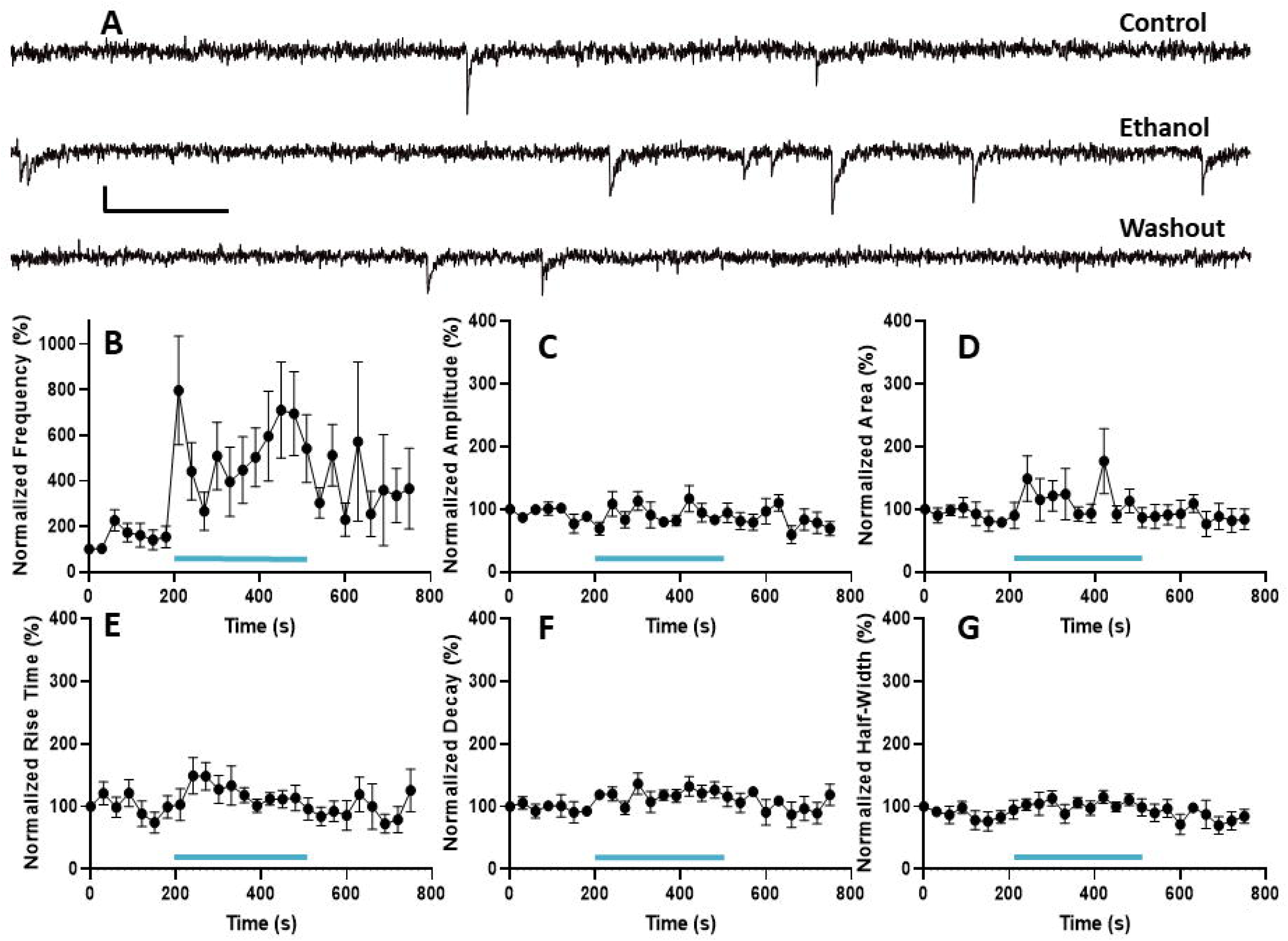
Acute ethanol exposure dose-dependently increases the frequency of action potential-dependent GABA_A_ receptor-mediated spontaneous postsynaptic currents (GABA_A_-sPSCs) in granule cells at postnatal days 7-10. A) Representative GABA_A_-sPSC traces from a granule cell recording obtained in the absence of ethanol (control), during application of ethanol (50 mM), and during washout. Scale bar is 307 ms × 40 pA. Mean effect of ethanol on the following GABA_A_-mPSC parameters: B) frequency, C) amplitude, D) area, E) rise time 10-90%, F) decay time, and G) half-width. Data were normalized to the zero-time point. The light blue line represents the timing of ethanol application during the slice electrophysiological recording. Mixed effects ANOVA found significant effects of ethanol on frequency (time, p=0.0094).

## Discussion

The findings of this study advance understanding of ethanol’s acute effects on GoC function in the developing mouse cerebellum. We show that developing GoCs express low levels of the Cl⁻ exporter KCC2, likely leading to depolarizing GABA_A_ receptor-mediated currents in some cells. Acute ethanol exposure dose-dependently increased the frequency of action potential-dependent GABA_A_-sPSCs in GoCs, without affecting action potential-independent GABA_A_-mPSCs. Ethanol produced variable effects on GoC spontaneous firing but more consistently increased GABA_A_-sPSC frequency in their target cells, the GrCs. These results reveal novel mechanisms that may underlie cerebellar dysfunction in FASD.

In the developing cerebellum of VGAT-Venus mice, we observed that KCC2 is predominantly expressed in the soma of Purkinje neurons, with lower expression levels in the molecular layer interneuron and GrC layers, findings consistent with a previous study in rats [35]. High-resolution confocal microscopy revealed that developing GoCs express low levels of KCC2 protein. To our knowledge, only one study has examined KCC2 expression in GoCs, and it focused on mature cells. Kozareva et al. [15] used single-cell RNA sequencing in adult mice to show that GoCs display variable expression of the *SLC12A5* gene, which encodes the KCC2 transporter, including a subset with minimal or no expression (see, https://singlecell.broadinstitute.org/single_cell/study/SCP795/). These observations indicate that GABA_A_ receptor activation may retain excitatory influence in a specific subpopulation of mature GoCs. Follow-up studies should explore GABA_A_ receptor function in GoCs across developmental stages, ideally using methods that preserve intracellular chloride gradients, such as gramicidin perforated-patch recordings. Additionally, single-cell RNA sequencing or *in situ* proteomics could further elucidate KCC2 expression dynamics in developing GoCs.

Acute ethanol exposure increased the frequency of action potential-dependent GABA_A_-sPSCs in developing GoCs, but only at higher concentrations (50 and 100 mM, equivalent to 230 and 460 mg/dL, respectively). No effect was observed at 25 mM (115 mg/dL), a concentration near the legal intoxication threshold (17.4 mM ≈ 80 mg/dL). Ethanol did not alter GABA_A_-sPSC parameters reflecting postsynaptic receptor function, including amplitude, area, rise time (10–90%), decay, or half-width. Similarly, 50 mM ethanol had no significant effect on any properties of action potential-independent GABA_A_-mPSCs. These findings suggest that acute ethanol selectively enhances action potential-dependent GABA release onto developing GoCs. If a comparable effect occurs in humans, it would likely require heavy, binge-level ethanol exposure during the third trimester of pregnancy. The increase in GABA_A_-sPSC frequency implies enhanced spontaneous firing of upstream GABAergic neurons at this developmental stage. While the identity of GABAergic inputs to developing GoCs remains unknown, data from adult mice suggest they may include Lugaro cells, other GoCs, and/or deep cerebellar nuclei inhibitory neurons [7–11]. Future studies should investigate how ethanol affects the excitability of these upstream GABAergic populations during development.

Loose-patch cell-attached recordings revealed that acute ethanol exposure has a variable effect on the firing activity of developing GoCs, with some exhibiting partially reversible potentiation, while others show no change. These findings suggest that a subpopulation of developing GoCs is sensitive to ethanol-induced enhancement of spontaneous action potential firing. Previous studies from several laboratories, including our own, have independently demonstrated that acute ethanol exposure increases spontaneous firing of mature GoCs in rats and mice, both *in vivo* and *in vitro* [19, 21, 22, 24–28]. The underlying mechanism in mature GoCs involves inhibition of neuronal nitric oxide synthase and its downstream targets—the Na^+^/K^+^ ATPase and K^+^ channels [21, 25, 26]. Whether a similar mechanism operates in developing GoCs remains unknown. It is also unclear whether variability in neuronal nitric oxide synthase activity and its downstream signaling pathways contributes to the heterogeneous ethanol-induced potentiation observed in these immature neurons. Additionally, the ethanol-induced increase of GABA_A_-sPSC frequency may partly drive enhanced excitability in a subset of developing GoCs, an effect potentially modulated by the level of KCC2 expression in these cells.

The increase in spontaneous action potential firing of developing GoCs was associated with enhanced spontaneous GABA_A_ receptor-mediated synaptic transmission in developing GrCs. This effect parallels published findings in mature GrCs from both rats and mice [21, 22, 27]. In developing GrCs, however, this increase in GABAergic input is likely to have an excitatory effect in a subset of cells expressing low levels of KCC2. Such excitatory GABAergic signaling could activate voltage-gated calcium channels, triggering Ca^2+^-dependent intracellular pathways and gene expression changes that may alter the normal developmental trajectory of these neurons. Moreover, ethanol-induced depolarization via GABA_A_ receptors could lead to excessive activation of NMDA receptors, potentially resulting in excitotoxicity. These mechanisms may contribute to the apoptotic response observed in GrCs following acute, binge-like ethanol exposure during the third-trimester-equivalent developmental period [20, 36–42].

In conclusion, our findings demonstrate that acute ethanol exposure enhances GABAergic transmission in developing GoCs and GrCs in the mouse cerebellum. The excitatory effects of GABA in subsets of developing neurons with low KCC2 expression may contribute to ethanol-induced disruptions in cerebellar circuit maturation, potentially underlying the neurodevelopmental deficits observed in FASD. This study offers new insight into the cellular mechanisms that could underlie ethanol’s neurotoxicity during cerebellar development. Future research should focus on how ethanol-induced disruptions in GoC and GrC synaptic transmission during development may interfere with the proper maturation of cerebellar circuits.

## Supporting information

Supplementary data tables (statistics)

## Acknowledgements

Supported by R21-AA030639 NIH grant. This research was also partially supported by UNM Comprehensive Cancer Center Support Grant NCI P30CA118100 and made use of the Fluorescence Microscopy and Cell Imaging shared resource. The slide scanner at the Center for Brain Recovery and Repair is supported by grant P20GM109089. Grammarly and ChatGPT were used to improve language and readability while preparing this work. After using these tools, the authors reviewed and edited the content as needed and take full responsibility for the publication’s content. The authors report there are no competing interests to declare.

## Author Contributions

Karick Jotty-Arroyo (data curation, formal analysis, investigation, writing - review & editing), Andrea Iturralde-Carrillo (data curation, formal analysis, investigation, writing - original draft and review & editing), Michael Paffett (data curation, formal analysis, investigation, writing - original draft and review & editing), Rafael Mancero-Montalvo (formal analysis, writing - review & editing), and C. Fernando Valenzuela (conceptualization; data curation; formal analysis; funding acquisition; project administration; supervision; writing - original draft and review & editing).

## References

1. Galliano, E., P. Mazzarello, and E. D’Angelo, Discovery and rediscoveries of Golgi cells. J Physiol, 2010. 588(Pt 19): p. 3639–55.

2. D’Angelo, E., et al., The cerebellar Golgi cell and spatiotemporal organization of granular layer activity. Front Neural Circuits, 2013. 7: p. 93.

3. Eccles, J.C., R. Llinas, and K. Sasaki, The mossy fibre-granule cell relay of the cerebellum and its inhibitory control by Golgi cells. Exp Brain Res, 1966. 1(1): p. 82–101.

4. Nietz, A.K., et al., Non-synaptic signaling from cerebellar climbing fibers modulates Golgi cell activity. Elife, 2017. 6.

5. Dumoulin, A., A. Triller, and S. Dieudonne, IPSC kinetics at identified GABAergic and mixed GABAergic and glycinergic synapses onto cerebellar Golgi cells. J Neurosci, 2001. 21(16): p. 6045–57.

6. Dieudonne, S. and A. Dumoulin, Serotonin-driven long-range inhibitory connections in the cerebellar cortex. J Neurosci, 2000. 20(5): p. 1837–48.

7. Szoboszlay, M., et al., Functional Properties of Dendritic Gap Junctions in Cerebellar Golgi Cells. Neuron, 2016. 90(5): p. 1043–56.

8. Hull, C. and W.G. Regehr, Identification of an inhibitory circuit that regulates cerebellar Golgi cell activity. Neuron, 2012. 73(1): p. 149–58.

9. Dugue, G.P., et al., Electrical coupling mediates tunable low-frequency oscillations and resonance in the cerebellar Golgi cell network. Neuron, 2009. 61(1): p. 126–39.

10. Ankri, L., et al., A novel inhibitory nucleo-cortical circuit controls cerebellar Golgi cell activity. Elife, 2015. 4.

11. Eyre, M.D. and Z. Nusser, Only a Minority of the Inhibitory Inputs to Cerebellar Golgi Cells Originates from Local GABAergic Cells. eNeuro, 2016. 3(2).

12. Schilling, K., Revisiting the development of cerebellar inhibitory interneurons in the light of single-cell genetic analyses. Histochem Cell Biol, 2024. 161(1): p. 5–27.

13. Hoshino, M., Molecular machinery governing GABAergic neuron specification in the cerebellum. Cerebellum, 2006. 5(3): p. 193–8.

14. Simat, M., et al., GABAergic synaptogenesis marks the onset of differentiation of basket and stellate cells in mouse cerebellum. Eur J Neurosci, 2007. 26(8): p. 2239–56.

15. Kozareva, V., et al., A transcriptomic atlas of mouse cerebellar cortex comprehensively defines cell types. Nature, 2021. 598(7879): p. 214–219.

16. Cancedda, L., et al., Excitatory GABA action is essential for morphological maturation of cortical neurons in vivo. J Neurosci, 2007. 27(19): p. 5224–35.

17. Ben-Ari, Y., Excitatory actions of gaba during development: the nature of the nurture. Nat Rev Neurosci, 2002. 3(9): p. 728–39.

18. Ouardouz, M. and B.R. Sastry, Activity-mediated shift in reversal potential of GABA-ergic synaptic currents in immature neurons. Brain Res Dev Brain Res, 2005. 160(1): p. 78–84.

19. Diaz, M.R., et al., Repeated intermittent alcohol exposure during the third trimester-equivalent increases expression of the GABA(A) receptor delta subunit in cerebellar granule neurons and delays motor development in rats. Neuropharmacology, 2014. 79: p. 262–74.

20. Backman, C., et al., Electrophysiological characterization of cerebellar neurons from adult rats exposed to ethanol during development. Alcohol Clin Exp Res, 1998. 22(5): p. 1137–45.

21. Kaplan, J.S., C. Mohr, and D.J. Rossi, Opposite actions of alcohol on tonic GABA(A) receptor currents mediated by nNOS and PKC activity. Nat Neurosci, 2013. 16(12): p. 1783–93.

22. Santhakumar, V., et al., A reinforcing circuit action of extrasynaptic GABAA receptor modulators on cerebellar granule cell inhibition. PLoS One, 2013. 8(8): p. e72976.

23. Diaz, M.R., et al., Na+/K+-ATPase inhibition partially mimics the ethanol-induced increase of the Golgi cell-dependent component of the tonic GABAergic current in rat cerebellar granule cells. PLoS One, 2013. 8(1): p. e55673.

24. Huang, J.J., et al., Acute ethanol exposure increases firing and induces oscillations in cerebellar Golgi cells of freely moving rats. Alcohol Clin Exp Res, 2012. 36(12): p. 2110–6.

25. Botta, P., et al., Excitation of rat cerebellar Golgi cells by ethanol: further characterization of the mechanism. Alcohol Clin Exp Res, 2012. 36(4): p. 616–24.

26. Botta, P., et al., Alcohol excites cerebellar Golgi cells by inhibiting the Na+/K+ ATPase. Neuropsychopharmacology, 2010. 35(9): p. 1984–96.

27. Carta, M., M. Mameli, and C.F. Valenzuela, Alcohol enhances GABAergic transmission to cerebellar granule cells via an increase in Golgi cell excitability. J Neurosci, 2004. 24(15): p. 3746–51.

28. Freund, R.K., Y. Wang, and M.R. Palmer, Differential effects of ethanol on the firing rates of Golgi-like neurons and Purkinje neurons in cerebellar slices in vitro. Neurosci Lett, 1993. 164(1-2): p. 9–12.

29. Wang, Y., et al., Fluorescent labeling of both GABAergic and glycinergic neurons in vesicular GABA transporter (VGAT)-venus transgenic mouse. Neuroscience, 2009. 164(3): p. 1031–43.

30. Nagai, T., et al., A variant of yellow fluorescent protein with fast and efficient maturation for cell-biological applications. Nat Biotechnol, 2002. 20(1): p. 87–90.

31. Bankhead, P., et al., QuPath: Open source software for digital pathology image analysis. Sci Rep, 2017. 7(1): p. 16878.

32. Schneider, C.A., W.S. Rasband, and K.W. Eliceiri, NIH Image to ImageJ: 25 years of image analysis. Nat Methods, 2012. 9(7): p. 671–5.

33. Saha, B., et al., Interactomic analysis reveals a homeostatic role for the HIV restriction factor TRIM5alpha in mitophagy. Cell Rep, 2022. 39(6): p. 110797.

34. Noureddine, A., et al., Endolysosomal Mesoporous Silica Nanoparticle Trafficking along Microtubular Highways. Pharmaceutics, 2021. 14(1).

35. Mikawa, S., et al., Developmental changes in KCC1, KCC2 and NKCC1 mRNAs in the rat cerebellum. Brain Res Dev Brain Res, 2002. 136(2): p. 93–100.

36. Nirgudkar, P., et al., Ethanol exposure during development reduces GABAergic/glycinergic neuron numbers and lobule volumes in the mouse cerebellar vermis. Neurosci Lett, 2016. 632: p. 86–91.

37. Bonthius, D.J., Jr., et al., Importance of genetics in fetal alcohol effects: null mutation of the nNOS gene worsens alcohol-induced cerebellar neuronal losses and behavioral deficits. Neurotoxicology, 2015. 46: p. 60–72.

38. Heaton, M.B., et al., Differential effects of ethanol on c-jun N-terminal kinase, 14-3-3 proteins, and Bax in postnatal day 4 and postnatal day 7 rat cerebellum. Brain Res, 2012. 1432: p. 15–27.

39. Botia, B., et al., Neuroprotective effects of PACAP against ethanol-induced toxicity in the developing rat cerebellum. Neurotox Res, 2011. 19(3): p. 423–34.

40. Nowoslawski, L., B.J. Klocke, and K.A. Roth, Molecular regulation of acute ethanol-induced neuron apoptosis. J Neuropathol Exp Neurol, 2005. 64(6): p. 490–7.

41. Luo, J., J.R. West, and N.J. Pantazis, Nerve growth factor and basic fibroblast growth factor protect rat cerebellar granule cells in culture against ethanol-induced cell death. Alcohol Clin Exp Res, 1997. 21(6): p. 1108–20.

42. Hamre, K.M. and J.R. West, The effects of the timing of ethanol exposure during the brain growth spurt on the number of cerebellar Purkinje and granule cell nuclear profiles. Alcohol Clin Exp Res, 1993. 17(3): p. 610–22.

